# Associations between volumes of grey matter regions and white matter hyperintensities with cognitive empathy in oldest-old adults

**DOI:** 10.1101/2025.10.16.682959

**Authors:** Russell J. Chander, Sarah A. Grainger, John D. Crawford, Jiyang Jiang, Nicole A. Kochan, Katya Numbers, Henry Brodaty, Julie D. Henry, Wei Wen, Perminder S Sachdev

## Abstract

**OBJECTIVES:** To study the differential associations of regional volumes of grey matter (GM) and white matter hyperintensities (WMH) with behavioural and informant-reported measures of cognitive empathy (CE), especially in brain regions understood to have functional connectivity involvement with CE abilities.

**METHODS:** Community-dwelling participants from the Sydney Memory and Ageing Study (Sydney MAS) underwent whole brain MRI. Regional cortical GM volumes were derived using FreeSurfer v7.1 for, and WMH volumes were derived using UBO Detector. On follow-up four years later, CE was indexed via the Reading the Mind in the Eyes Test (RMET; a behavioural task) and Interpersonal Reactivity Index – Perspective Taking subscale (IRI-PT; an informant-reported measure).

**RESULTS:** In 129 participants (mean age 87.01 years), Structural equation modelling (SEM) showed associations between insula GM volumes and RMET scores (p=0.001), and between supramarginal gyrus (SMG) GM volumes and IRI-PT scores (p=0.016). Associations remained significant after inclusion of covariates accounting for age, global cognition, affective symptoms, and social networks.

**DISCUSSION:** CE abilities assessed with RMET (a behavioural task) were positively associated with insula GM volumes. CE abilities on the IRI-PT (an informant-reported measure) were positively associated with SMG GM volumes. No associations between WMH volumes and CE measures were found. These findings may inform clinical workflows that use structural MRI investigations looking at social cognitive disturbances in older adults and provide further evidence of the divergent nature of behavioural and self-report measures of CE.

## INTRODUCTION

### Cognitive empathy in older age

Social cognition refers to the cognitive processes that people engage to perceive and interpret social cues in their environment, which helps to inform social judgment and behaviour (Henry et al., 2015). Theory of mind, affective empathy, social perception, and social behaviour have been identified as the four core domains of social cognitive function (Henry et al., 2016). Social cognitive failures across one or multiple domains is an important feature of many neurodegenerative diseases, and is a component criterion of Version 5 of the Diagnostic and Statistical Manual (DSM-5) for neurocognitive disorders, alongside other more traditionally recognized subdomains such as memory and executive function (American Psychiatric Association, 2013). Extensive literature now also shows that social cognitive difficulties are often seen in the context of normal adult ageing (Beadle & de la Vega, 2019; Hayes et al., 2020; Henry et al., 2013) with potential implications for broader mental health and wellbeing (Kelly et al., 2017; Masi et al., 2011). Several mechanisms have been proposed to explain social cognitive change in late adulthood (Grainger et al., 2024; Henry et al., 2023; Kynast et al., 2018), one of which is age-related brain changes, particularly in the default mode network (Hughes et al., 2019; Moran et al., 2012).

The aspect of social cognition of particular interest to the present study is cognitive empathy (CE), which is closely aligned with the construct of affective theory of mind and refers to our understanding of others’ emotions, affective states, or feelings (Premack & Premack, 1995; Rogers et al., 2007). This is understood to be distinct from affective empathy, which refers to our emotional response to the perceived situations of others and how these emotions may be similar or distinct to the emotions of the other (Mehrabian & Epstein, 1972).

There has been some work done looking at the functional activity of specific brain regions under CE tasks in functional magnetic resonance imaging (fMRI) studies. CE has been linked to activity in various brain regions, such as the medial prefrontal cortex (mPFC) (Adolphs, 2001; Moran et al., 2012; Suzuki et al., 2019), insula (Krendl & Kensinger, 2016; Riva et al., 2018), and the anterior cingulate cortex (ACC) (Krendl & Kensinger, 2016). Certain brain regions are also implicated in multiple social cognitive skills, and are thus not specialized to CE. An example of this is the insula, where activity has been associated with the specific CE ability to use context to inform feelings of pity (Krendl & Kensinger, 2016). Decreased activity levels in the anterior insula have also been observed in older adults actively empathizing with targets of different emotional expressions (Riva et al., 2018), which is more akin to affective empathy.

### Limitations of existing literature

There is significant heterogeneity in the literature with regards to which brain regions have been shown to be associated with CE (Krendl & Kensinger, 2016; Riva et al., 2018; Suzuki et al., 2019; Ziaei et al., 2016). One possible explanation for this lies in the various CE measures used, which may have introduced some variability in how CE is indexed. Prior work has utilised bespoke experimental paradigms (Krendl & Kensinger, 2016; Riva et al., 2018; Suzuki et al., 2019; Ziaei et al., 2016), self-report scales (Riva et al., 2018), and behavioural or psychometric assessments (Ziaei et al., 2016). While these methods have been validated for CE, there have been noted differences in how older adults perform in certain social cognition tasks, and CE tasks have been especially noted to have poor correlation between each other (Grainger et al., 2023; Murphy & Lilienfeld, 2019). Certain behavioural or self-report CE tasks may also be more salient depending on the nature of the cohort studied and level of neurocognitive impairment expected. For example, prior work indicates that behavioural assessments of CE are more associated with early neurocognitive changes seen in mild cognitive impairment, while informant-rated changes might indicate more significant changes that are mainly seen in people with dementia (Chander et al., 2024). This makes it all the more likely that behavioural and informant-rated CE measures are related but diverging constructs of CE with distinct underlying neurological substrates. Since the majority of existing studies have looked at the association between brain regions and CE performance as indexed by one measure (for example, see work on the relationship between a behavioural CE task and the right middle temporal gyrus (Otsuka et al., 2024)), more evidence is needed to investigate how performance in multiple types of CE measures might be associated with different brain regions.

Another gap in the literature lies in how prior research has mainly focused on neural activation and task-based fMRI paradigms. While such work is highly informative, especially in looking for early subtle brain changes, there remains a need to understand what structural brain changes are associated with empathy in older adults. Structural MRI studies remain crucial to the clinical diagnostic process, as a great degree of neuroimaging investigations in psychiatric and neurological clinical settings in older adults utilize structural MRI scans in their primary investigations. Furthermore, functional connectivity between regions as observed in fMRI are at times linked to subtle structural changes in these regions (Meyer-Lindenberg et al., 2005). Research knowledge that identifies key brain volumes associated with empathic deficits in older adults can be readily implemented in clinical investigations. This also needs to be considered given that there is already some level of brain changes that are expected in the normal ageing process (Fox & Schott, 2004). Understanding what brain changes are associated with healthy ageing and with empathic deficits can help us understand if these changes are themselves also part of the expected ageing process, or indicative of neurocognitive disorders such as dementia.

Additionally, there is a paucity of research looking at the effects of white matter hyperintensities (WMH) on social skills and social cognitive abilities in older adults. WMH lesions detected in fluid-attenuated inversion recovery (FLAIR) MRI scans are indicative of cerebrovascular disease (CeVD), and generally accumulate in older adult brains (Wardlaw et al., 2013). CeVD had been associated with impaired cerebral blood flow and functional connectivity (Huang et al., 2021), and generalized burden of CeVD by total volume of WMH has been associated with poorer cognitive and social cognitive outcomes (Kynast et al., 2018). Etiological differences between regional manifestations of WMH have been noted, with periventricular WMH (pvWMH) being more associated with cognitive impairment and deep WMH (dWMH) with mood disorders in older adults (Kim et al., 2008). More information is needed on if pvWMH or dWMH lesions would differentially have an impact on CE performance.

*Study objective*

The primary objective of the current study was to identify key brain imaging regions that are associated with behavioural and self-reported measures of CE performance in older adults, and to investigate how CeVD possibly impacts on CE. We hypothesized that:

1. Performance in behavioural and informant-rated measures of CE is associated with volumes in differential cortical grey matter regions, including the frontal and limbic cortices,
2. Performance in behavioural and informant-rated measures of CE is differentially associated with volumes of pvWMH and dWMH.

## METHODS

### Cohort and recruitment

Participants were selected from the Sydney Memory and Ageing Study (Sydney MAS) cohort, the details of which have been previously published (Sachdev et al., 2010). Briefly, 1,037 community participants aged 65 years and older were recruited from two electoral districts within Sydney, New South Wales, Australia between 2005 and 2007. At recruitment, inclusion criteria included at least basic literacy and conversational English, and that participants do not have any major neurological, psychological, or psychiatric diagnoses. At baseline (Wave 1), and then subsequently every two years in Waves 2-6, participants underwent a comprehensive assessment that included neuropsychological assessments, questionnaires about psychological wellbeing and quality of life, physical assessments, and medical background assessments. Baseline interviews also included the collection of demographic information, including if participants came from a non-English speaking background (NESB). Participants also underwent additional measures at certain Waves, such as MRI studies. Study partners of participants, who were commonly spouses, offspring, or close relatives of the participants, were also invited to fill out questionnaires about participants’ everyday function and psychological well-being.

The current study focuses on data collected in Waves 4 and 6, which occurred six and ten years after Baseline respectively. Wave 4 included an MRI study and was the most recent Wave to include such protocols. Wave 6 was the first Wave in Sydney MAS to include social cognition measures. Thus, data from these two Waves offered the most contemporaneous neuroimaging and social cognition data available in Sydney MAS. At Wave 4, 258 participants consented to an additional neuroimaging study and had completed MRI protocols that passed quality control for use in volumetric analysis. At Wave 6, 441 participants provided at least some social cognition measurement data. An additional 10 participants were excluded for having Mini-Mental State Examination (MMSE)(Folstein et al., 1975) scores at Wave 6 below 24, to exclude possible dementia newly identified at this Wave. From the overlap of the two data sources, the sample size of the current study is 129 participants with both Wave 4 MRI data and Wave 6 social cognition data. Waves 4 and 6 of Sydney MAS will subsequently be referred to as Timepoint 1 (T1) and Timepoint 2 (T2) for this study.

The Sydney MAS study was conducted in accordance with institutional ethical guidelines and approved by the UNSW Human Research Ethics Committee (HREC; HC: 05037, 09382, 14327, 190962). All participants and their informants provided voluntary informed consent prior to their involvement in the Sydney MAS study.

### Neuroimaging protocol

At T1, participants were invited to participate in an additional MRI study. Scans were performed on a Philips 3T Achieva Quasar Dual scanner. Acquisition parameters for T1-weighted structural MRI scans were: TR = 6.39 ms, TE = 2.9 ms, flip angle = 8°, matrix size = 256 × 256, FOV = 256 × 256 × 190, and slice thickness = 1 mm with no gap in between, yielding 1 × 1 × 1 mm^3^ isotropic voxels.

Raw imaging files were processed in FreeSurfer v7.1.0 (http://surfer.nmr.mgh.harvard.edu/) to obtain data on cortical structures using Desikan-Killiany Atlas. Briefly, the processing steps included skull stripping, Tailarach registration, spherical surface maps, and parcellation of cortex which consisted of distinct cortical regions, including 13 frontal, nine temporal, four occipital, seven parietal, and the insula for each hemisphere as per the Desikan-Killiany atlas (Desikan et al., 2006). In order to control for individual intracranial volumes (ICV) in our analysis, cortical volumes were expressed as percentages of the estimated total ICV estimated by FreeSurfer.

WMH volumes were derived using UBO Detector, an automated pipeline that uses T1-weighted and FLAIR MRI images to classify, segment, and measure WMH volumes based on anatomical location, intensity levels, and cluster size (Jiang et al., 2018). T1-weighted and FLAIR images were brought into DARTEL space, where they were co-registered using the T1-weighted images as reference to reslice the FLAIR images in SPM12. T1 images were then used to generate probability maps for GM, WM, and CSF, and the probability maps were brought to DARTEL space along with the T1 and coregistered FLAIR images. Non-brain elements were then removed and FMRIB’s FAST segmentation was used to identify WMH clusters (Zhang et al., 2001). A k-nearest neighbours algorithm (Warfield et al., 2000) was then used to classify WMH volumes, and the derived volumes were further parcellated into periventricular (pvWMH) and deep (dWMH), and these values were re-expressed as percentages of total ICV values derived as above. Further details on the UBO Detector pipeline have been previously published (Jiang et al., 2018).

### ROI selection

In an effort to manage the number of studied variables, a literature search was conducted to identify previous work that highlighted specific brain regions of interest (ROIs) as having a relationship with social cognition. Papers were selected to inform the model based on the following criteria: 1) at least one aspect of empathy was measured, and 2) structural or functional neuroimaging was conducted.

In order to capture as many empathy studies as possible, search parameters included alternative terms for cognitive empathy, such as mindreading, theory of mind, and intention attribution.

Regions that did not directly align with the regions in the current data were reconciled. The most common discrepancy was the literature identifying larger brain regions where the atlas instead identified the constituent regions. For example, the prefrontal cortex (PFC) was identified by multiple publications as a region of interest in CE (Adolphs, 2001; Frith & Frith, 2006; Gallagher et al., 2000; Moran et al., 2012; Suzuki et al., 2019; Ziaei et al., 2016). While the Desikan-Killany atlas did not specifically identify the PFC, it did identify the lateral and medial orbitofrontal cortices (OFCs), and the caudal and rostral anterior cingulate cortices (ACCs). This was resolved by combining the lateral and medial OFC into a combined OFC region, and ACC volumes were similarly generated. Lateral volumetric variables were also totalled together to generate one bilateral volume for each ROI. Supplementary Table 1 also includes the constituent brain volumes identified by the atlas that were combined to tally with brain regions in the literature.

### Cognitive empathy measures

In addition to clinical and neuropsychological assessments at T2, participants completed several social cognitive assessments. Of these measurements, the Reading the Mind in the Eyes Test (RMET)(Baron-Cohen et al., 2001) and the Interpersonal Reactivity Index (IRI)(Davis, 1980) were of particular interest here. The RMET is a 36-item behavioural measure of CE that asks participants to identify what a presented target is most likely to be thinking based on only their eyes, out of four possible options. A total score out of 36 is generated based on how many correct responses a participant has. Of the current study sample, 116 participants have complete RMET and neuroimaging data.

The IRI is an informant-rated scale that asks the study partner to rate the participant’s performance on a series of empathy scenarios based on a 5-point Likert scale. The specific subscale used in this study was the 7-item Perspective Taking (IRI-PT) subscale for CE, which includes items such as asking how likely the participant is to adopt the other person’s point of view when in a disagreement. Of the current study sample, 66 participants have complete IRI-PT and neuroimaging data.

Besides social cognition measures, additional assessments were conducted at T2 as part of the standard Sydney MAS study visit. General cognitive ability was measured using the MMSE (Folstein et al., 1975). The 15-item Geriatric Depression Scale (GDS)(Sheikh & Yesavage, 1986) was used to assess depressive symptoms, and the Goldberg Anxiety Scale (GAS)(Goldberg et al., 1988) was used to index symptoms of anxiety. Additionally, the Lubben Social Network Scale (LSNS)(Lubben et al., 2006) was used to provide a measure of the extent of the participant’s social network of friends and family. These measures were identified for inclusion in the current study due to their potential confounding effects on social cognition.

### Statistical analysis

Linear regression models were run to select the neuroimaging variables most associated with CE scores. Two models were run with RMET scores and IRI-PT scores from T2 as the outcome respectively, and all GM and WMH volume variables of interest as predictors, along with age and scores for MMSE, GDS, GAS, and LSNS as covariates. Reverse stepwise elimination of variables was run at a p <0.10 threshold to remove nonsignificant variables. This was done to identify the unique neuroimaging variables most associated with RMET scores and IRI-PT scores individually.

Structural equation modelling (SEM) via the *lavaan* package (Rosseel, 2012) was then used to investigate the relationships between neuroimaging volumes and CE measures while also accounting for confounding variables in the models. Two separate models were run for RMET scores (Model 1; Figure 1) and IRI-PT scores (Model 2; Figure 2), and each model included the GM and/or WMH variables of interest as identified in linear regression models, age at T1 and T2, MMSE at T2, GDS scores at T2, and a ‘Social networks’ latent factor derived from LSNS Friends and Family subscales scores. The regressions of all GM and/or WMH variables of interest on CE measures were modelled, along with the effect of T1 age on all neuroimaging variables, T2 age on CE measures, MMSE scores, the ‘Affective’ factor, and the ‘Social network’ factor, the effect of T2 MMSE scores on CE measures, and covariance between T1 and T2 age. Earlier iterations of the model that fit GAS and GAS scores onto an ‘Affective’ latent factor produced coefficients greater than 1, likely indicating Heywood cases. Since GAS scores did not significantly converge onto the ‘Affective’ latent factor, GAS scores were dropped from the models, and GDS scores were studied as an observed factor. Coefficients greater than 1 were similarly observed with the regression of LSNS Friend scores on the ‘Social network’ factor. This was resolved by fixing this specific path coefficients to 0.999.

**Figure 1:**
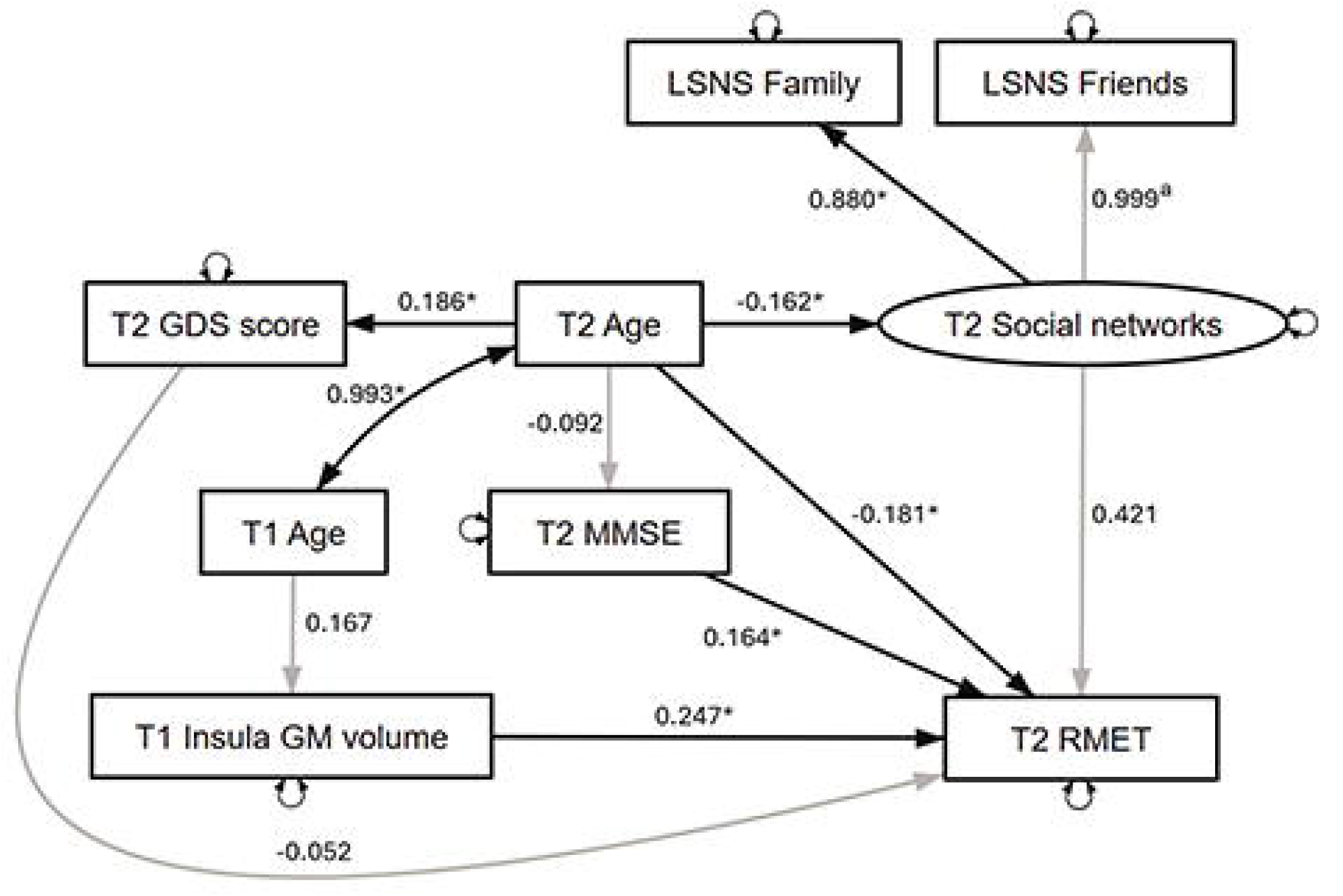
SEM model predicting T2 RMET scores. Paths in bold indicate statistically significant regressions and covariances, and values indicate standardized coefficients. Abbreviations: GDS, Geriatric Depression Scale; GM, grey matter; LSNS, Lubben Social Network Scale; MMSE, Mini-Mental Scale Examination; RMET, Reading the Mind in the Eyes Test; T1, Timepoint 1 (aligning to Wave 4 of Sydney MAS); T2, Timepoint 2 (aligning to Wave 6 of Sydney MAS). * p < 0.05 ^a^ path coefficient locked to 0.999 to address Heywood cases.

**Figure 2:**
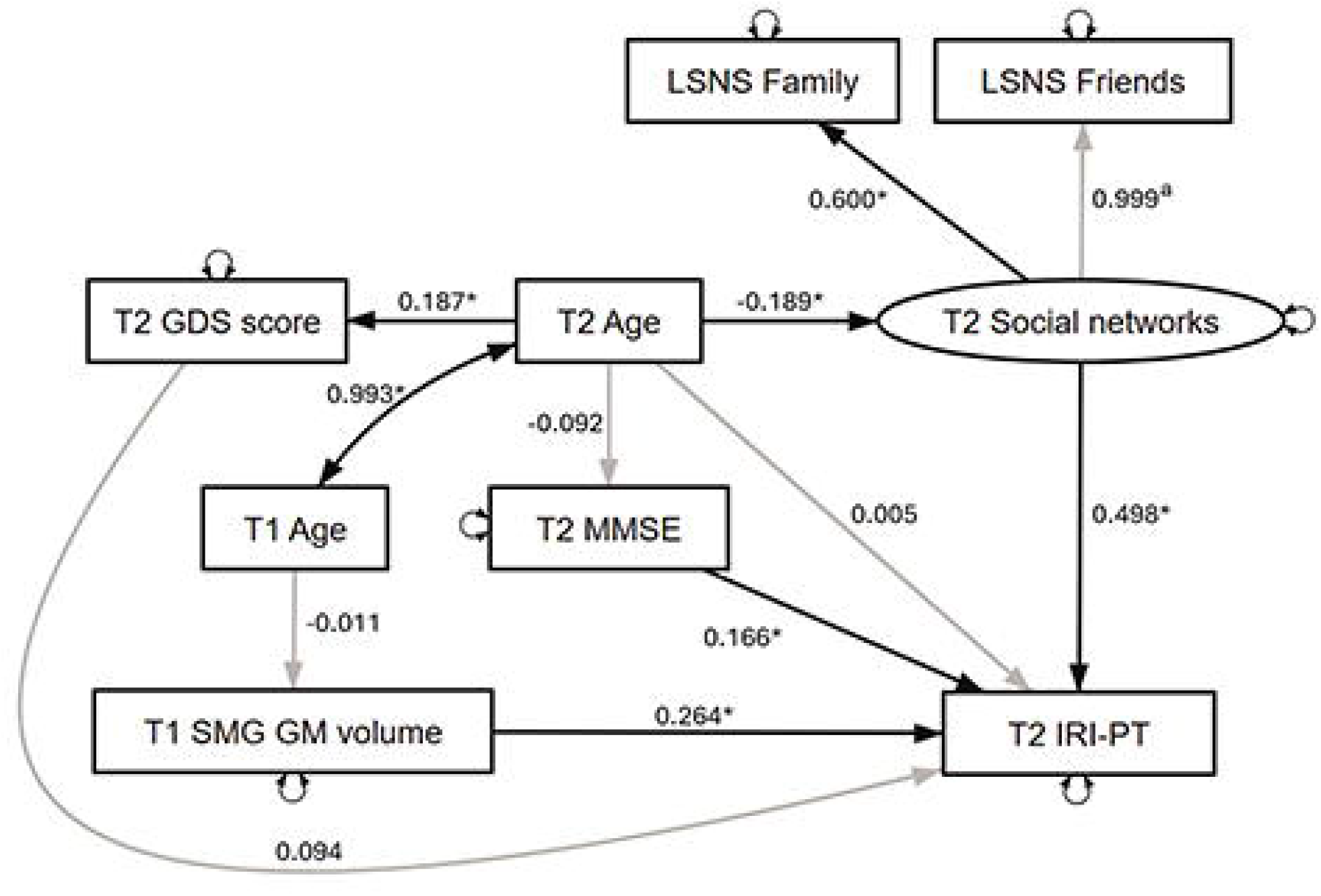
SEM model predicting T2 IRI-PT scores. Paths in bold indicate statistically significant regressions and covariances, and values indicate standardized coefficients. Abbreviations: GDS, Geriatric Depression Scale; GM, grey matter; IRI-PT, Interpersonal Reactivity index – Perspective Taking subscale; LSNS, Lubben Social Network Scale; MMSE, Mini-Mental Scale Examination; SMG, supramarginal gyrus; T1, Timepoint 1 (aligning to Wave 4 of Sydney MAS); T2, Timepoint 2 (aligning to Wave 6 of Sydney MAS). * p < 0.05 ^a^ path coefficient locked to 0.999 to address Heywood cases.

Descriptive analyses and regression models were conducted in Stata version 15.1 (StataCorp LLC, College Station, TX). SEM analyses were conducted in R v3.6.0 (R Core Team, 2019), and the SEM model figures were drawn using the open source *semdiag* package (Mai et al., 2023). All p-value thresholds were set at <0.05, except for stepwise elimination of variables in linear regression models, where a less conservative threshold of p <0.10 was used.

## RESULTS

### Demographics

A total of 170 participants were included in the current study. At T2, these participants had a mean age of 87.01 years (SD 4.13 years), with an interquartile range of 83.45 – 89.65 years, and a min-max range of 80.72 – 100.08 years. 97.7% of participants were Caucasian, 56.5% were female, and 12.9% were NESB (Table 1). Compared to the 867 participants who were not included in the current study, study participants that were included were younger at the original baseline visit of Sydney MAS, and there was a lower proportion of MCI participants at original baseline. Included and excluded participants did not differ significantly in terms of gender, NESB status, or educational attainment (Supplementary Table 2).

**Table 1:** Descriptive data for T1 neuroimaging data and T2 assessment results (N = 129). Reported values for the RMET and IRI-PT were for the subset of participants that had this data (n = 119 for RMET, n = 69 for IRI-PT).

| Demographics |  |
| --- | --- |
| Age at T2, mean (SD), y | 86.58 (3.88) |
| Age at T2, IQR, y | 83.29 – 89.27 |
| Gender, N (%), female | 81 (62.8%) |
| Ethnicity, N (%), Caucasian | 126 (97.7%) |
| Education, mean (SD), y | 12.05 (3.68) |
| NESB, N (%) | 14 (10.9%) |
| Time in study as of T2, mean (SD), y | 10.37 (0.16) |
| Time in study as of T2, min-max | 10.08 – 10.91 |
| T1 Neuroimaging |  |
| Orbitofrontal cortex, mean (SD), % of eTIV | 1.65 (0.17) |
| Precuneus, mean (SD), % of eTIV | 1.20 (0.13) |
| Insula, mean (SD), % of eTIV | 0.12 (0.10) |
| Anterior cingulate cortex, mean (SD), % of eTIV | 0.53 (0.08) |
| Supramarginal gyrus, mean (SD), % of eTIV | 1.26 (0.16) |
| dWMH, mean (SD), % of eTIV | 0.49 (0.62) |
| pvWMH, mean (SD), % of eTIV | 1.01 (0.68) |
| T2 Assessments |  |
| MMSE score, mean (SD) | 28.98 (1.32) |
| RMET score, mean (SD) | 22.00 (4.93) |
| IRI-PT, mean (SD) | 23.13 (5.69) |
| GDS score (depression), mean (SD) | 2.21 (2.03) |
| GAS score (anxiety), mean (SD) | 2.16 (2.09) |
| LSNS family score, mean (SD) | 23.87 (5.77) |
| LSNS friends score, mean (SD) | 20.77 (5.69) |

**Notes:** dWMH, deep white matter hyperintensities; eTIV, estimated total intracranial volume; GAS, Goldberg Anxiety Scale; GDS, Geriatric Depression Scale; IQR, interquartile range; IRI-PT, Interpersonal Reactivity Index – Perspective Taking subscale; LSNS, Lubben Social Network Scale; MMSE, Mini-Mental State Examination; pvWMH, periventricular white matter hyperintensities; RMET, Reading the Mind in the Eyes Test.

At T2, participants scored an average of 28.31 points on the MMSE. 119 participants had RMET data, and 69 participants had IRI data (Table 1). Consistent with prior work (Grainger et al., 2023; Murphy & Lilienfeld, 2019), behavioural (i.e. RMET) and self-report (i.e. IRI-PT) scores were not correlated with each other (Pearson’s r = 0.13, p = 0.305). Therefore, these two variables were not loaded onto a single latent CE factor but treated as separate endogenous variables in the linear regression and SEM analyses.

### ROI identification and volumes

Based on the literature search, the following brain regions were identified as key ROIs: OFC, precuneus, insula, ACC, and supramarginal gyrus (SMG) (Supplementary Table 1). Additionally, to examine the effects of WMH on social cognition, pvWMH and dWMH volumes were also included (Table 1).

Linear regression models using neuroimaging variables to predict CE outcomes were run (Table 2). The model with RMET scores as the outcome found that, while no individual neuroimaging measure was a significant predictor, insula GM volumes were trending towards significance. After reverse stepwise elimination of variables, insula GM volumes remained as the sole significant neuroimaging predictor of RMET scores (β = 0.22, 95% CI 0.04 – 0.40; p = 0.017). The model with IRI-PT scores as the outcome found that SMG GM volumes were significantly associated with IRI-PT scores out of all the neuroimaging variables and also survived stepwise elimination of variables (β = 0.23, 95% CI 0.00 – 0.45; p = 0.053). Neither pvWMH nor dWMH volumes were significant in either regression model. Based on this, insula GM volumes were included in the SEM analysis predicting RMET scores, and SMG GM volumes were included in the SEM analysis predicting IRI-PT scores.

**Table 2:** Linear regression models of neuroimaging variables and covariates predicting RMET (n = 119) and IRI-PT (n = 69) scores before and after stepwise elimination.

| Outcome: T2 RMET score | Full model |  | After stepwise elimination |  |
| --- | --- | --- | --- | --- |
| | $\beta$ (95% CI) | p value | $\beta$ (95% CI) | p value |
| OFC GM volume | -0.11 (-0.40 – 0.17) | 0.433 | – | – |
| ACC GM volume | 0.15 (-0.07 – 0.37) | 0.186 | – | – |
| Precuneus GM volume | -0.18 (-0.44 – 0.08) | 0.172 | – | – |
| Insula GM volume | 0.23 (-0.04 – 0.49) | 0.094 | 0.22 (0.04 – 0.40) | 0.017* |
| SMG GM volume | 0.06 (-0.18 – 0.30) | 0.608 | – | – |
| pvWMH volume | -0.03 (-0.36 – 0.30) | 0.849 | – | – |
| dWMH volume | 0.08 (-0.24 – 0.40) | 0.634 | – | – |
| Age | -0.23 (-0.43 – -0.03) | 0.024* | -0.23 (-0.41 – -0.05) | 0.013* |
| MMSE score | 0.21 (0.03 – 0.40) | 0.025* | 0.15 (-0.03 – 0.33) | 0.095 |
| GDS score | 0.03 (-0.18 – 0.24) | 0.780 | – | – |
| GAS score | 0.12 (-0.09 – 0.34) | 0.264 | – | – |
| LSNS Family score | 0.04 (-0.14 – 0.23) | 0.635 | – | – |
| LSNS Friend score | 0.19 (0.00 – 0.38) | 0.049 | – | – |

| Outcome: T2 IRI-PT score | Full model |  | After stepwise elimination |  |
| --- | --- | --- | --- | --- |
| | $\beta$ (95% CI) | p value | $\beta$ (95% CI) | p value |
| OFC GM volume | -0.13 (-0.56 – 0.29) | 0.526 | – | – |
| ACC GM volume | -0.13 (-0.44 – 0.18) | 0.399 | – | – |
| Precuneus GM volume | -0.20 (-0.57 – 0.16) | 0.271 | – | – |
| Insula GM volume | 0.14 (-0.29 – 0.57) | 0.519 | – | – |
| SMG GM volume | 0.43 (0.06 – 0.81) | 0.025* | 0.23 (0.00 – 0.45) | 0.053 |
| pvWMH volume | 0.38 (-0.10 – 0.85) | 0.117 | – | – |
| dWMH volume | -0.24 (-0.68 – 0.18) | 0.253 | – | – |
| Age | -0.22 (-0.53 – 0.08) | 0.147 | -0.21 (-0.45 – 0.02) | 0.076 |
| MMSE score | 0.10 (-0.15 – 0.35) | 0.415 | – | – |
| GDS score | 0.11 (-0.21 – 0.44) | 0.488 | – | – |
| GAS score | -0.20 (-0.50 – 0.11) | 0.204 | – | – |
| LSNS Family score | 0.17 (-0.10 – 0.43) | 0.212 | 0.25 (0.01 – 0.49) | 0.039* |
| LSNS Friend score | 0.20 (-0.07 – 0.47) | 0.140 | – | – |

**Notes:** ACC, anterior cingulate cortex; dWMH, deep white matter hyperintensities; GAS, Goldberg Anxiety Scale; GDS, Geriatric Depression Scale; GM, grey matter; IRI-PT, Interpersonal Reactivity Index – Perspective Taking subscale; LSNS, Lubben Social Network Scale; MMSE, Mini-Mental State Examination; OFC, orbitofrontal cortex; pvWMH, periventricular white matter hyperintensities; RMET, Reading the Mind in the Eyes Test; SMG, supramarginal gyrus; T2, study timepoint 2.
\* p < 0.05

### SEM analysis

SEM Model 1 for RMET scores converged after 39 iterations and exhibited excellent fit [comparative fit index (CFI): 0.933; Tucker-Lewis index (TLI): 0.887; root mean square error of approximation (RMSEA) = 0.034]. The regression of insula GM volumes to RMET scores was significant (p=0.001), indicating that higher RMET scores were associated with greater insula GM volumes. Other significant predictors for RMET score included T2 age and T2 MMSE scores (Figure 1, Supplementary Table 3).

SEM Model 2 for IRI-PT scores converged after 41 iterations and also exhibited excellent fit [CFI: 0.997; TLI: 0.960; RMSEA = 0.017]. The regression of SMG GM volumes to IRI-PT scores was significant (p=0.016), indicating that higher IRI-PT scores were associated with greater SMG GM volumes. In this model, T2 MMSE scores and the ‘Social network’ factor were the only other significant predictor of IRI-PT scores (Figure 2, Supplementary Table 4).

## DISCUSSION

This study explores the association between structural brain regions and CE skills four years later in a cohort of older community-dwelling adults via SEM. The hypothesis that performance in CE measures was differentially associated with specific frontal and limbic grey matter volumes was partially supported. Overall, RMET (a behavioural CE task) was positively associated with insula GM volumes, and IRI-PT (an informant-reported CE measure) was positively associated with SMG GM volumes. These associations remained significant after accounting for effects of age, cognition, affective symptoms, and social connectedness. The hypothesis that performance in CE measures was differentially associated with pvWMH or dWMH volumes was not supported, as WMH volumes were not found to be associated with either CE measure.

The finding that larger insula volumes are associated with better behavioural CE scores is corroborated in previous work linking insula activation and empathic attitudes (Krendl & Kensinger, 2016; Ziaei et al., 2016). One study found that global cognition was a factor in older adults’ ability to express empathic attitudes to outgroup members in emergency situations. In addition, this study also revealed that insula activation was associated with how malleable the individual’s attitudes were to contextual environmental information regarding how responsible the outgroup member was for their own plight (Krendl & Kensinger, 2016). Considering how CE includes skills in reading people’s intentions and drawing inferences from contextual cues, it would follow that the insula is associated with CE specifically via the ability to consolidate social cues and information.

The finding of SMG GM volumes being associated with informant-reported CE measures is corroborated by previous work finding that lower neural activation in the right SMG is associated with a greater degree of emotional egocentricity, which refers to the tendency of one’s current emotional state to bias one’s ability to exercise empathy towards another (Riva et al., 2016). The SMG is a region of the brain that tends to experience age-related decline from early in the lifespan (Sowell et al., 2003), which would indicate that such age-related changes to the SMG, and thus CE skills, are expected to a certain degree.

Despite the fact that the RMET and IRI-PT are both measures of CE, these two measures are likely indexing different components of CE. This has been shown in prior work that demonstrated how performance in behavioural tasks and self or informant reports of CE are unrelated (Grainger et al., 2023). This potentially explains the differing brain regions associated with RMET and IRI-PT scores in this study. Furthermore, IRI-PT was significantly associated with MMSE scores, but these scores had no significant relationship with age, which suggests that these observed MMSE changes were not age-related and might indicate early neurocognitive impairments. It is possible that insula GM changes are associated with early CE changes, while SMG GM changes are associated with more generalized and marked CE deficits that might be related to early neurocognitive disorders. This would indicate some potential in paying specific attention to insula and SMG GM volumes to indicate early brain changes that might herald future conversion to dementia.

It should be noted that all selected ROIs have been identified in prior work examining age-related empathic abilities (see work discussing the mPFC (Moran et al., 2012; Suzuki et al., 2019), precuneus (Suzuki et al., 2019), and ACC (Krendl & Kensinger, 2016)). However, most regions were not found to be significant in the current analysis. Most prior work has focused on functional activation levels within these regions, as opposed to the volumetric approach taken in the current analysis. It has been suggested that structural integrity and functional activity are not always correlated in the brain (Schulz et al., 2022). While functional changes may help index one way in which the brain undergoes early changes that reflect in cognitive abilities, structural changes might be a more prominent marker of neural degradation that warrants attention.

Despite previous work finding a negative association between WMH volumes and social cognition (Kynast et al., 2018), the current analysis did not find a relationship between WMH (as total volume or periventricular/deep volumes) and measures of CE. It should be noted that compared to participants in the current study, the cohort studied by Kynast and colleagues were significantly younger, with a mean age of 60 years (SD 13.1) and a range or 21-79 years. This study also found that having a Fazekas score of 3 was associated with poorer social cognition scores, which represents severe and significant WMH burden. Such a high amount of WMH burden would be atypical, especially in that age range, and this is reflected in how Fazekas 3 participants made up just under 2% of their cohort. This might suggest that these individuals were experiencing a high degree of cerebrovascular burden associated with significant cognitive disruptions and neurocognitive disorders. In the current old-old cohort free of major neurocognitive disorders, the observed burden of WMH might not have had as great of an effect on CE and other cognitive and social cognitive skills as other age-related changes, such as GM volumes, would have.

The current findings have some applicability in a clinical setting. In the same vein as early detection and prevention of cognitive decline, methods that aid in early detection of possible social cognitive change can aid clinical management workflows in identifying at-risk individuals, with the intention of more closely monitoring such individuals for future change. Brain changes that are associated with age-related social cognitive changes can also serve as additional information when distinguishing between mild and pathological social cognitive changes, such as those seen in late-life psychiatric conditions.

This study uniquely explored structural brain volumes associated with both behavioural and informant-rated measures of CE abilities in healthy older adults. The advanced age and large sample size of the cohort are also strengths as this age group is particularly understudied. Some limitations of the current work should be addressed. Due to the availability of data and protocols of Sydney MAS, non-contemporaneous neuroimaging and social cognition data that was 4 years apart had to be used. However, given the relatively high MMSE scores and inclusion of nondemented older adults only, this cohort is unlikely to have experienced significant neurodegeneration. Thus, there is a high degree of confidence that the brain volumes derived from T1 are likely to be stable and similar by T2. Nevertheless, there are still restrictions to the kind of conclusions that can be derived from the current work.

In conclusion, cognitive empathy performance in older adults is significantly associated with differential GM volumes, specifically the insula for behavioural CE scores and the SMG for informant-rated CE measures. Further work is needed to investigate longitudinal changes in brain volumes and social cognitive performance to establish a directional relationship, as well as to determine if degeneration in specific brain regions is associated with distinct levels of social cognitive performance.

## AUTHOR CONTRIBUTIONS

R.J.C. contributed to the study concept and design, interpretation of the data, and drafting of the manuscript. S.A.G. contributed to the interpretation of the data and drafting of the manuscript. J.D.C contributed to the statistical design of the study. J.J. and W.W. contributed to the acquisition, processing, and interpretation of the neuroimaging data. K.N. contributed to data acquisition. N.A.K. and H.B. contributed to cohort design of Sydney MAS. J.D.H. contributed to study concept and design, interpretation of the data, and drafting of the manuscript. P.S.S. contributed to the study concept and design, and cohort design of Sydney MAS. All authors contributed to the editing of the manuscript for intellectual content.

## DATA STATEMENT

A preliminary version of this analysis was submitted as part of a thesis in fulfilment of the requirements for the Doctor of Philosophy for R.J.C. Data and statistical code pertaining to this study are available upon request.

## ACKNOWLEDGMENTS

The authors would like to thank all current and former study team members of Sydney MAS for their contributions and guidance towards the study, as well as all study participants for volunteering their time and effort to be a part of Sydney MAS. We would also like to thank Dr. Ben Lam for his statistical guidance on this analysis.

