## Supplementary Table 1 for "Associations between volumes of grey matter regions and white matter hyperintensities with cognitive empathy in oldest-old adults"

**Supplementary Table 1:** Key regions of interest identified as being associated with cognitive and affective empathy, and variable matching with ROIs in the Desikan-Killany atlas.

| **Brain region**  **(References)** | **Corresponding region(s) in atlas** | **Social cognition subdomains affected** | **Variable management notes** |
| --- | --- | --- | --- |
| Medial prefrontal cortex (1–6) | Lateral orbitofrontal  Medial orbitofrontal  Caudal anterior cingulate  Rostral anterior cingulate  Frontal pole | Theory of mind / Cognitive empathy (1–6) | Combine lateral and medial orbitofrontal regions into orbitofrontal cortex  Combine caudal and rostral anterior cingulate cortex regions into anterior cingulate cortex |
| Precuneus (1) | Precuneus | Theory of mind / Cognitive empathy (1) | None |
| Insula (4,7,8) | Insula | Facial emotion recognition (4)  Affective empathy (7,8) | None |
| Anterior cingulate cortex (4,7,9,10) | Caudal anterior cingulate  Rostral anterior cingulate | Facial emotion recognition (4)  Affective empathy (7,10)  Theory of mind / Cognitive empathy (9) | Combine caudal and rostral anterior cingulate regions into anterior cingulate cortex |
| Supramarginal gyrus (11) | Supramarginal | Theory of mind / Cognitive empathy (11) | None |
