## Supplementary Table 2 for "Associations between volumes of grey matter regions and white matter hyperintensities with cognitive empathy in oldest-old adults"

**Supplementary Table 2:** Comparisons between participants included in current study and excluded prior to study.

|  | **Included in study**  **N = 170** | **Excluded from study N = 867** | **p value** |
| --- | --- | --- | --- |
| Wave 1 age, mean (SD), years | 76.16 (4.12) | 78.76 (4.83) | <0.001* |
| Wave 1 age, med (IQR), years | 76 (73 - 79) | 79 (75 - 82) | - |
| Wave 1 age, min - max, years | 70 - 89 | 70 - 90 | - |
| Gender, no. (%), female | 96 (56.5%) | 476 (54.9%) | 0.707 |
| NESB, no. (%) | 22 (12.9%) | 142 (16.4%) | 0.261 |
| Education, mean (SD), years | 11.90 (3.58) | 11.54 (3.45) | 0.149 |
| Wave 1 MCI status, no. (%) | 47 (30.5%) | 286 (39.1%) | 0.046* |

**Abbreviations:** MCI, mild cognitive impairment; NESB, non-English speaking background.
