## Supplementary Table 3 for "Associations between volumes of grey matter regions and white matter hyperintensities with cognitive empathy in oldest-old adults"

**Supplementary Table 3:** Structural equation modelling results for Model 1 predicting RMET score.

|  | **β** | **Standard error** | **95% confidence interval** | **p value** |
| --- | --- | --- | --- | --- |
| **Latent Variables** | | | | |
| Social 🡨 |  |  |  |  |
| LSNS Family score | 0.880 | 0.435 | 0.028 – 1.731 | 0.043 |
| LSNS Friend score | 0.999 | - | - | - |
| **Regressions** | | | | |
| T2 RMET score 🡨 |  |  |  |  |
| T1 Insula GM volume | 0.247 | 0.075 | 0.100 – 0.394 | 0.001 |
| T2 Age | -0.181 | 0.063 | -0.304 – -0.058 | 0.004 |
| T2 MMSE score | 0.164 | 0.051 | 0.064 – 0.264 | 0.001 |
| T2 GDS score | -0.052 | 0.054 | -0.158 – 0.054 | 0.336 |
| Social | 0.421 | 0.297 | -0.162 – 1.004 | 0.157 |
| T1 Insula GM vol 🡨 |  |  |  |  |
| T1 Age | 0.167 | 0.086 | -0.002 – 0.336 | 0.053 |
| T2 MMSE score 🡨 |  |  |  |  |
| T2 Age | -0.092 | 0.059 | -0.208 – 0.025 | 0.122 |
| T2 GDS score 🡨 |  |  |  |  |
| T2 Age | 0.186 | 0.055 | 0.078 – 0.295 | 0.001 |
| Social 🡨 |  |  |  |  |
| T2 Age | -0.162 | 0.073 | -0.306 – -0.018 | 0.027 |
| **Covariances** | | | | |
| T1 Age 🡨🡪 T2 Age | 0.993 | 0.066 | 0.864 – 1.123 | <0.001 |
| **Intercepts** | | | | |
| LSNS Family score | -0.005 | 0.055 | -0.113 – 0.102 | 0.922 |
| LSNS Friend score | -0.002 | 0.054 | -0.108 – 0.103 | 0.963 |
| T2 RMET score | -0.010 | 0.054 | -0.116 – 0.096 | 0.851 |
| T1 Insula GM volume | 0.020 | 0.086 | -0.149 – 0.188 | 0.818 |
| T2 MMSE score | 0.000 | 0.054 | -0.105 – 0.105 | >0.999 |
| T2 GDS score | 0.002 | 0.054 | -0.103 – 0.107 | 0.965 |
| T2 Age | 0.000 | 0.054 | -0.106 – 0.106 | >0.999 |
| T1 Age | 0.000 | 0.054 | -0.106 – 0.106 | >0.999 |
| **Variances** | | | | |
| LSNS Family score | 0.765 | 0.146 | 0.479 – 1.052 | <0.001 |
| LSNS Friend score | 0.696 | 0.162 | 0.379 – 1.013 | <0.001 |
| T2 RMET score | 0.800 | 0.066 | 0.670 – 0.930 | <0.001 |
| T1 Insula GM volume | 0.966 | 0.127 | 0.717 – 1.215 | <0.001 |
| T2 MMSE score | 0.989 | 0.097 | 0.799 – 1.179 | <0.001 |
| T2 GDS score | 0.962 | 0.106 | 0.755 – 1.169 | <0.001 |
| Social | 0.275 | 0.136 | 0.009 – 0.541 | 0.042 |
| T2 Age | 0.997 | 0.066 | 0.868 – 1.126 | <0.001 |
| T1 Age | 0.997 | 0.066 | 0.867 – 1.127 | <0.001 |

**Abbreviations:** GDS, Geriatric Depression Scale; GM, grey matter; LSNS, Lubben Social Network Scale; MMSE, Mini-Mental State Examination; RMET, Reading the Mind in the Eyes Test; T1, study timepoint 1; T2, study timepoint 2.
