## Supplementary Table 4 for "Associations between volumes of grey matter regions and white matter hyperintensities with cognitive empathy in oldest-old adults"

**Supplementary Table 4:** Structural equation modelling results for Model 2 predicting IRI-PT score.

|  | **β** | **Standard error** | **95% confidence interval** | **p value** |
| --- | --- | --- | --- | --- |
| **Latent Variables** | | | | |
| Social 🡨 |  |  |  |  |
| LSNS Family score | 0.600 | 0.244 | 0.122 – 1.078 | 0.014 |
| LSNS Friend score | 0.999 | - | - | - |
| **Regressions** | | | | |
| T2 IRI-PT score 🡨 |  |  |  |  |
| T1 SMG GM volume | 0.264 | 0.110 | 0.049 – 0.480 | 0.016 |
| T2 Age | 0.005 | 0.081 | -0.154 – 0.163 | 0.955 |
| T2 MMSE score | 0.166 | 0.065 | 0.040 – 0.293 | 0.010 |
| T2 GDS score | 0.094 | 0.082 | -0.067 – 0.254 | 0.253 |
| Social | 0.498 | 0.252 | 0.004 – 0.992 | 0.048 |
| T1 SMG GM vol 🡨 | ~ |  |  |  |
| T1 Age | -0.011 | 0.085 | -0.178 – 0.155 | 0.894 |
| T2 MMSE score 🡨 |  |  |  |  |
| T2 Age | -0.092 | 0.059 | -0.208 – 0.025 | 0.122 |
| T2 GDS score 🡨 |  |  |  |  |
| T2 Age | 0.187 | 0.055 | 0.078 – 0.295 | 0.001 |
| Social 🡨 |  |  |  |  |
| T2 Age | -0.189 | 0.057 | -0.301 – -0.078 | 0.001 |
| **Covariances** | | | | |
| T1 Age 🡨🡪 T2 Age | 0.993 | 0.066 | 0.864 – 1.123 | <0.001 |
| **Intercepts** | | | | |
| LSNS Family score | -0.004 | 0.055 | -0.111 – 0.103 | 0.938 |
| LSNS Friend score | -0.003 | 0.054 | -0.108 – 0.102 | 0.962 |
| T2 IRI-PT score | 0.016 | 0.072 | -0.124 – 0.156 | 0.823 |
| T1 SMG GM volume | -0.007 | 0.086 | -0.177 – 0.162 | 0.934 |
| T2 MMSE score | 0.000 | 0.054 | -0.105 – 0.105 | >0.999 |
| T2 GDS score | 0.002 | 0.054 | -0.103 – 0.107 | 0.972 |
| T2 Age | 0.000 | 0.054 | -0.106 – 0.106 | >0.999 |
| T1 Age | 0.000 | 0.054 | -0.106 – 0.106 | >0.999 |
| **Variances** | | | | |
| LSNS Family score | 0.839 | 0.108 | 0.628 – 1.051 | <0.001 |
| LSNS Friend score | 0.557 | 0.188 | 0.188 – 0.925 | 0.003 |
| T2 IRI-PT score | 0.778 | 0.110 | 0.563 – 0.993 | <0.001 |
| T1 SMG GM volume | 0.991 | 0.129 | 0.739 – 1.244 | <0.001 |
| T2 MMSE score | 0.989 | 0.097 | 0.799 – 1.179 | <0.001 |
| T2 GDS score | 0.962 | 0.106 | 0.755 – 1.169 | <0.001 |
| Social | 0.405 | 0.178 | 0.055 – 0.754 | 0.023 |
| T2 Age | 0.997 | 0.066 | 0.868 – 1.126 | <0.001 |
| T1 Age | 0.997 | 0.066 | 0.867 – 1.127 | <0.001 |

**Abbreviations:** GDS, Geriatric Depression Scale; GM, grey matter; IRI-PT, Interpersonal Reactivity Index – Perspective Taking subscale; LSNS, Lubben Social Network Scale; MMSE, Mini-Mental State Examination; SMG, supramarginal gyrus; T1, study timepoint 1; T2, study timepoint 2.
